# The encapsulin from *Thermatoga maritima* is a flavoprotein with a symmetry matched ferritin-like cargo protein

**DOI:** 10.1101/2021.04.26.441214

**Authors:** Benjamin LaFrance, Caleb Cassidy-Amstutz, Robert J Nichols, Luke M Oltrogge, Eva Nogales, David F Savage

**Author notes:** these authors contributed equally (Benjamin LaFrance, Caleb Cassidy-Amstutz).

## Abstract

Bacterial nanocompartments, also known as encapsulins, are an emerging class of protein-based ‘organelles’ found in bacteria and archaea. Encapsulins are virus-like icosahedral particles comprising a ∼25-50 nm shell surrounding a specific cargo enzyme. Compartmentalization is thought to create a unique chemical environment to facilitate catalysis and isolate toxic intermediates. Many questions regarding nanocompartment structure-function remain unanswered, including how shell symmetry dictates cargo loading and to what extent the shell facilitates enzymatic activity. Here, we explore these questions using the model *T. maritima* nanocompartment known to encapsulate a redox-active ferritin-like protein. Biochemical analysis revealed the encapsulin shell to possess a flavin binding site located at the interface between capsomere subunits, suggesting the shell may play a direct and active role in the function of the encapsulated cargo. Furthermore, we used cryoEM to show that cargo proteins use a form of symmetry-matching to facilitate encapsulation and define stoichiometry. In the case of the *T. maritima* encapsulin, the decameric cargo protein with 5-fold symmetry preferentially binds to the pentameric-axis of the icosahedral shell. Taken together, these observations suggest the shell is not simply a passive barrier—it also plays a significant role in the structure and function of the cargo enzyme.

## Introduction

Subcellular compartmentalization is a common strategy to sequester metabolic pathways that may be incompatible with the wellbeing of the rest of the cell. Although compartmentalization was originally thought of as a hallmark feature of eukaryotes, many prokaryotes use proteinaceous compartments to facilitate biochemical reactions. For example, bacterial microcompartments, such as the carboxysome, are formed from hundreds to thousands of copies of numerous different subunits (*1*–*3*). A new family of bacterial protein compartments, known as encapsulins, are increasingly realized to also occur throughout prokaryotic phyla (*4*). Encapsulins possess a less complex architecture and form smaller compartments than their microcompartment counterparts and are therefore referred to as nanocompartments. Encapsulins are often composed of a single type of shell and cargo proteins and are usually limited to around 25-50 nm in diameter (*5, 6*). The cargo protein typically has enzymatic function and is often related to redox chemistry *(7, 8)*.

One example of the redox role within encapsulins is found in the hyperthermophilic bacterium *Thermatoga maritima*, where the cargo is a ferritin-like protein (FLP). FLPs function like ferritin to sequester soluble iron, Fe(II), from the intracellular environment by oxidation and mineralization in an Fe(III) form. Mineralization prevents soluble iron from spontaneously reacting with reactive oxygen species as demonstrated in the Fenton reaction, which can damage the cell. This was shown to be the case for *M. xanthus*, for which cells with a disrupted or deleted encapsulin gene showed significantly lower survival rates following hydrogen peroxide exposure than wildtype *M. xanthus* (*9*). Alternatively, He *et al*. demonstrated that FLP-loaded encapsulins can store nearly nine times more iron atoms than apoferritin, suggesting FLP-loaded encapsulins are a cellular iron mega-storage compartment (*10*).

The first extensive biochemical characterization of a nanocompartment was that of the *T. maritima* encapsulin by Sutter *et al* in 2008 (*4*). This sample was purified from the native host and x-ray crystallographic experiments resolved the 3.1Å structure of the encapsulin shell as an icosahedron built from 12 pentameric capsomeres (i.e. T=1 symmetry). Although the cargo was present in the sample, occupancy-related issues precluded determining the FLP cargo structure, save for a C-terminal eight amino-acid sequence that was identified near the 5-fold axis of the encapsulin. Follow-up work has shown this peptide is sufficient for cargo targeting (*11, 12*). Based on homology modeling and related structures, the FLP cargo was hypothesized to have 5-fold symmetry by Sutter, and later confirmed through structure determination by He *et al* (*10*). Because the cargo FLP symmetry is 5-fold, and there is also a 5-fold symmetry axis on the icosahedral shell, it was thought that there may be symmetry matching that occurs between encapsulin shells and their cargo proteins.

Some groups have characterized the degree of encapsulin loading for large cargo proteins (*4, 8*); however, the assumption for a small cargo like FLP is that every binding site is loaded. For FLP this would actually be a 2:1 ratio stoichiometry of FLP monomer:encapsulin protomer because the FLP is a 5-fold symmetric decamer that binds to the pentameric shell vertex (*4, 10*). For the *B. linens* DyP encapsulin, only a single cargo protein was encapsulated despite additional space within the shell (*13, 14*). However, a recent study of the *M. smegmatis* DyP encapsulin found both a minority species of the encapsulin contained a single hexameric DyP cargo while the majority of particles actually contained a dodecameric DyP complex from 2 DyP cargo interacting with one another (*15*). Furthermore, encapsulation of native and non-native cargo appears to cause significant destabilization of the nanocompartment (*13, 14*), which may explain the absence of higher cargo loading ratios in many of these systems. Although many attempts have been made to visualize a cargo protein in complex within an encapsulin shell protein, it is likely that the substoichiometry, heterogeneity, and flexibility of the encapsulin-cargo interaction have prevented high-resolution structure determination that accurately resolves the cargo (*4, 5, 9, 16*–*19*).

In order to obtain a higher ratio of FLP to encapsulin shell we designed an expression system to maximize FLP production relative to the shell protein. Characterization of this complex by cryoEM allowed us to successfully resolve the structure of the native cargo protein in complex with the shell protein after symmetry expansion and extensive focused classification. Although the resolution does not allow accurate atomic modelling, helices are defined and rigid-body docking confirms the symmetry matching of the decameric FLP with the 5-fold axis of the shell. Additionally, spectroscopic analysis as well as mass spectrometry confirmed that this *T. maritima* encapsulin is a flavoprotein that binds flavin moieties across the interface of two shell subunits. The high-resolution cryoEM structure of the shell revealed that the FMN interacts intimately with a tryptophan, W90, between two shell subunits. The work presented here accurately resolves an FLP cargo in relation to the encapsulin shell, and reveals that the *T. maritima* encapsulin is a flavoprotein.

## Results and Discussion

### The *T. maritima* encapsulin is a flavoprotein

During purification of the *T. maritima* encapsulin, fractions containing the encapsulin protein were found to possess a yellow coloration (Figure 1 A, B). Yellow coloration is typical of flavoproteins and suggested that the *T. maritima* encapsulin may be a flavoprotein (*20*). To test this idea, we compared the absorbance spectrum of purified encapsulins to flavin mononucleotide (FMN) and riboflavin. Both FMN and riboflavin have local maxima at ∼350 nm and a global maxima at ∼450 nm, absorption features that are also shared by the *T. maritima* encapsulin (Figure 1 C). Endogenous encapsulin purified by Sutter *et al*. from *T. maritima* also had a yellow coloration, suggesting our observation was not a feature of recombinant expression but a fundamental feature of *T. maritima* encapsulins (*4, 21*). This result strongly supports the claim that *T. maritima* encapsulins are flavoproteins. Although this is an unexpected finding it has been recently independently proposed from a concurrent structural study (*22*).

**Figure 1.**
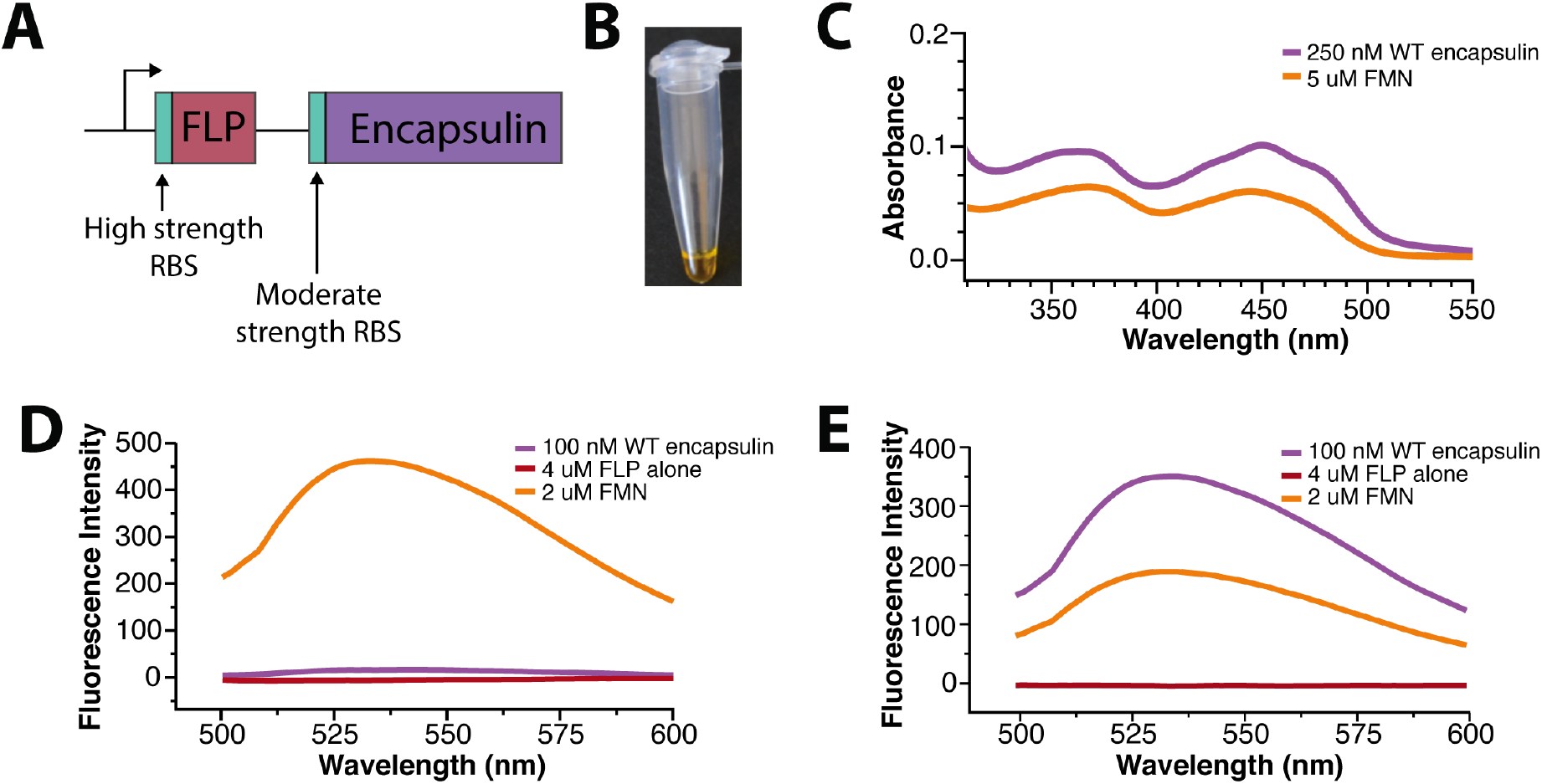
Spectroscopic characterization of the *T. maritima* encapsulin. **(A)** Strategy utilized to maximize cargo loading within the encapsulin shell. **(B)** *T. maritima* encapsulin exhibits a yellow hue upon purification. For panels C-E, the colors absorbance spectrum for WT encapsulin (FLP-cargo and shell) is purple, and FMN is orange, and the purified FLP cargo alone in maroon. **(C)** Encapsulin spectrum has maxima and minima similar to that of FMN. Fluorescence emission after excitation at 450nm for native samples **(D)** and under denaturing conditions **(E)**.

The intracellular flavin pool is composed of three major species that vary in size and redox potential—riboflavin, FMN, and flavin adenine dinucleotide (FAD). We sought to identify the specific cofactor bound to the encapsulin shell. As these three molecules are challenging to distinguish by UV-Vis absorbance data, we opted to use mass spectrometry for characterization. Both FMN and riboflavin were present as the major and minor peaks respectively, while no peak for FAD was observed (Supplemental Figure 1). Additionally, a lumichrome peak, a common flavin photodegradation product, was observed. Due to variations in ionization efficiencies, it is not possible to conclude the exact flavin composition. Despite the relatively large peak for lumichrome in the MS data, there clear density for the R-group beyond the flavin aromatic rings, and lumichrome peak is generated as an artifact of the mass spectrometry workflow (Supplemental Figure 1). Taken together, these data suggest that encapsulins preferentially bind FMN as well as riboflavin and further confirms that the *T. maritima* encapsulin purified herein is a flavoprotein.

While free, oxidized flavins have a well-characterized fluorescence profile with excitation at ∼450nm and emission at ∼530nm, purified *T. maritima* encapsulin displayed no such fluorescence (Figure 1 D). However, denaturation of the encapsulin sample in 7 M guanidinium chloride resulted in a canonical free flavin fluorescence profile (Figure 1 E). This suggests that the encapsulin quenches bound flavins. In other flavoproteins, fluorescence is often quenched through a π-stacking interaction with tyrosine, phenylalanine, or tryptophan (*23, 24*), suggesting that encapsulin flavin binding site possesses an aromatic residue.

### The Tryptophan at position 90 is necessary for flavin binding, despite poor conservation among all encapsulins

In order to better characterize the flavin-encapsulin interaction, we determined the cryoEM structure of the purified encapsulin complex at 3.3Å resolution (Figure 2 A, Supplemental Table 1). The overall architecture of the encapsulin shell was markedly similar to the previously determined structure (pdb:3DKT) (*4*). However, upon close examination, there is additional density near W90 that corresponds to a flavin undergoing π-stacking with the tryptophan (Figure 2 B). This same Tryptophan-flavin stacking interaction was also observed in a concurrent structure (*22*).

**Figure 2.**
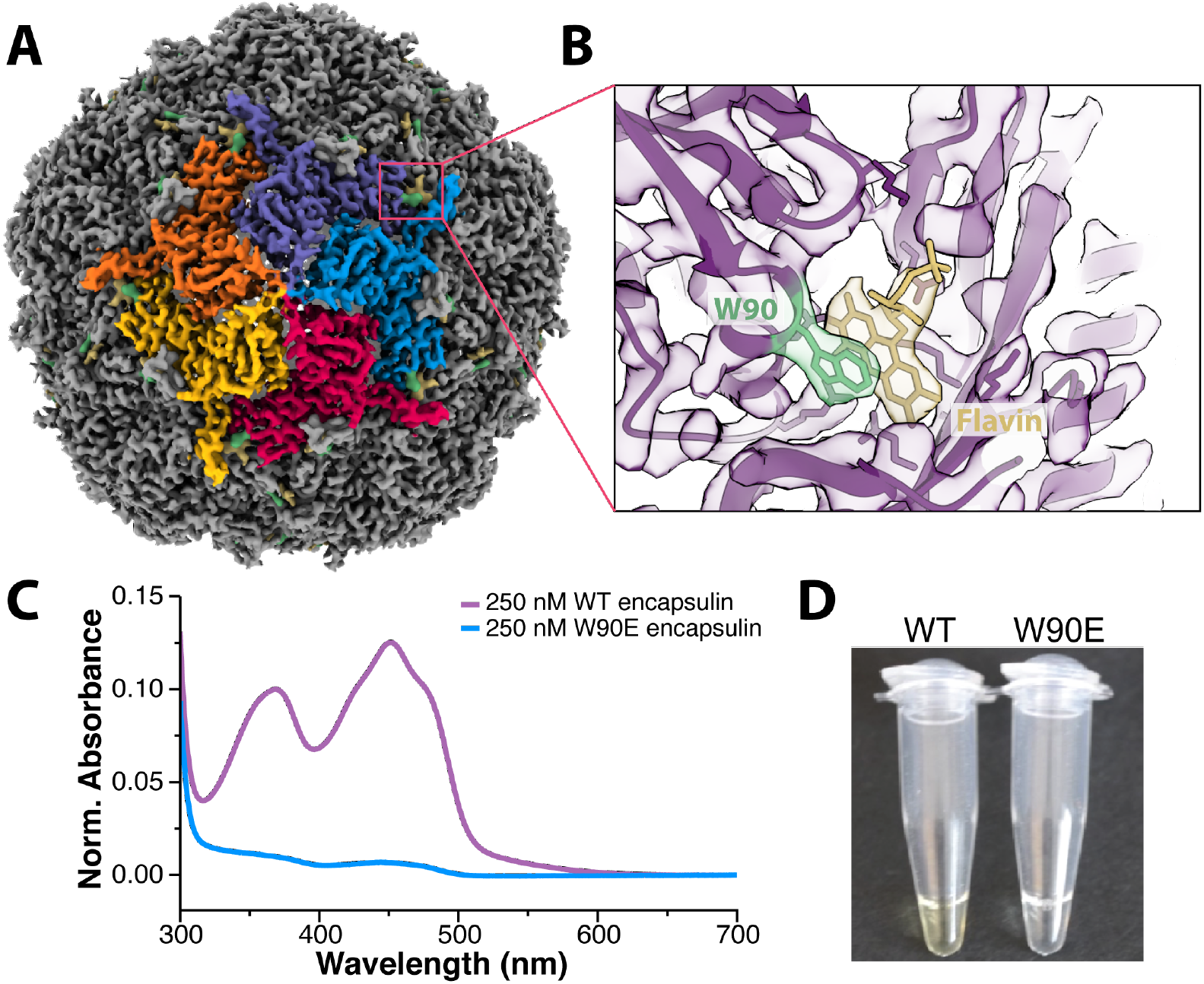
Structural determination confirms the presence of a flavin coordinated with W90. **(A)** CryoEM structure of the *T. maritima* encapsulin determined at 3.3Å resolution. **(B)** Zoomed-in view of a portion of the density map and corresponding atomic model showing the flavin density (in yellow) next to the tryptophan at position 90 (W90, in green). **(C)** Absorbance spectra of WT encapsulin (purple) and W90E (blue). **(D)** Color difference between wildtype and mutant W90E encapsulins after purification, protein concentration identical between the two samples.

To probe the importance of the W90 residue for flavin binding, a W90E mutation was made to remove the π-stacking interaction. The W90E mutant encapsulin purified similarly to wild-type *T. maritima* encapsulins. However, no yellow coloration was observed in the W90E sample during purification, and biochemical experiments revealed no flavin characteristics present for the W90E mutant encapsulin (Figure 2 C, D).

Having identified the flavin binding site for the *T. maritima* encapsulin, we sought to determine the potential of flavin binding for other homologs. Previous studies of other encapsulins have not noted yellow coloration of purified encapsulins, nor found flavins bound to the encapsulin shell. Interestingly, those encapsulins do not have an aromatic residue at position 90 (*4*–*7, 9*). Multiple sequence alignment revealed that most known encapsulins from other organisms do not share this critical tryptophan. In fact, a majority of known encapsulins do not contain an aromatic residue at this position. Of note, all encapsulins with an aromatic residue at position 90 also contain a ferritin-like protein (FLP) as their cargo. However, the inverse is not true and not all FLP encapsulins have the W90 residue and only 35% of known FLP-loaded encapsulins contain W90 (Supplemental Figure 2). Surprisingly, almost all of the FLP-loaded encapsulins with the W90 residue are from anaerobic organisms, suggesting flavin binding may only be necessary in anaerobic environments.

### The FLP cargo protein is flexibly bound within the encapsulin shell

Upon discovering that the *T. maritima* encapsulin was a flavoprotein, we next sought to better understand the role of the ferritin-like protein (FLP) cargo. In order to obtain a high-resolution structure of the encapsulin shell and fit the flavin molecule, icosahedral symmetry was imposed. However, imposing icosahedral symmetry reduced our ability to interpret the FLP cargo density, even though the cargo FLP was clearly visible in raw micrographs and 2D class averages (Figure 3 A, B). This could be due to sub-stoichiometric amounts of cargo loading (despite the over-expressions scheme in Figure 1 A), flexibility between the cargo FLP and the shell, or both.

**Figure 3.**
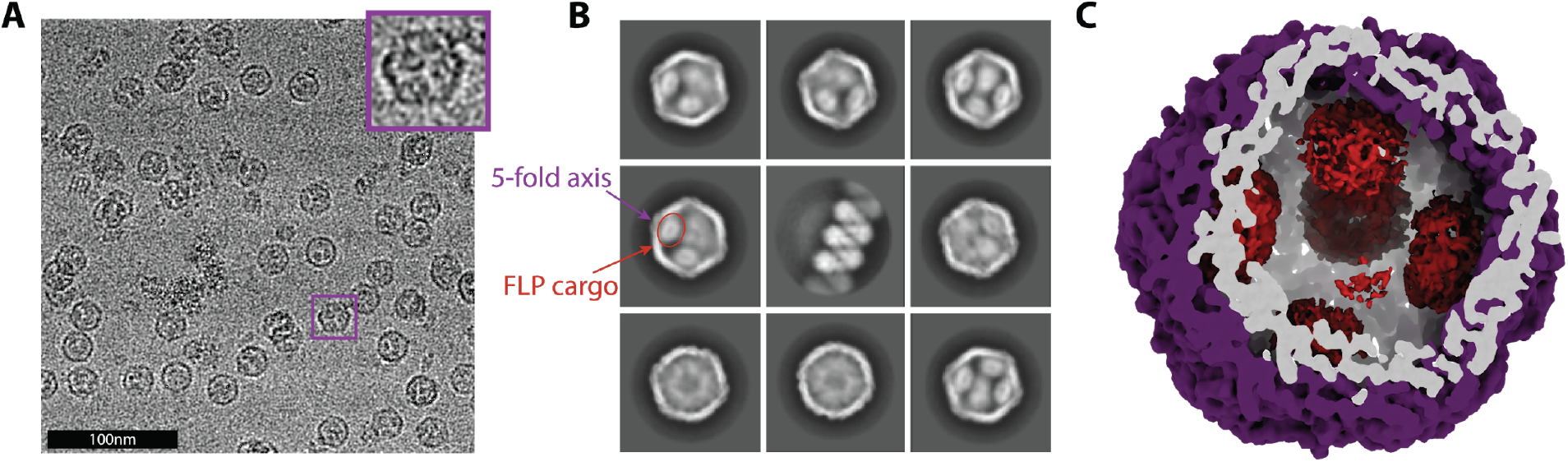
Initial cryo-EM visualization of the FLP-cargo protein within the encapsulin shell. **(A)** Raw micrograph. The inset shows an expanded view of a single particle to show that multiple FLP cargo proteins are clearly visible within the encapsulin, even before processing. **(B)** Curated 2D class averages further showing FLP density in some classes and illustrating differences in FLP loading: 2 FLPs (upper-left), 3-4 (bottom-right), as well as a small population lacking any cargo protein (bottom-left). **(C)** Slice through the encapsulin structure showing the shell in purple and weak, noisy FLP densities in red.

As an initial attempt to better visualize the cargo, we carried out an asymmetric C1 reconstruction. The C1 structure showed the presence of cargo, but the density was poorly defined (Figure 3 C). This result supports the idea of either sub-stoichiometric and inconsistent incorporation of cargo (i.e. some particles have 3 FLPs/shell while others have up to 5 FLPs/shell), or flexibility (i.e. each cargo FLP is not in the same exact location relative to the shell or the neighboring FLP). A small subset of particles appeared to contain 4 FLPs/shell in a tetrahedral style geometry as seen for the *H. ochraceum* FLP-encapsulin (*19*), but the density remained too weak to resolve accurately. In an attempt to improve the cargo density, signal subtraction was performed to remove signal from the encapsulin shell, which drives the alignment during the reconstruction. This procedure led to improved density for the cargo protein, but still not of enough resolution to identify the orientation or specific location of the cargo protein relative to the shell.

In order to resolve the structural heterogeneity of the FLP cargo within the encapsulin shell and better identify the interactions governing the FLP-encapsulin assembly, we applied symmetry expansion followed by a focused 3D classification strategy (Supplemental Figure 3, see Materials and Methods). First, we used the icosahedral symmetry of the encapsulin to place all 12 of the pentameric vertices of the shell—and their associated FLP cargo—into the same 3D location (Supplemental Figure 3). Next, we performed alignment-free 3D classification of that single location. This strategy significantly improved the resolution of the FLP (Figure 4 A). While still not high enough to build an atomic model, this resolution allowed us to rigid-body dock a crystal structure of a homologous decameric FLP previously reported (Figure 4 B,C) (*10*).

**Figure 4.**
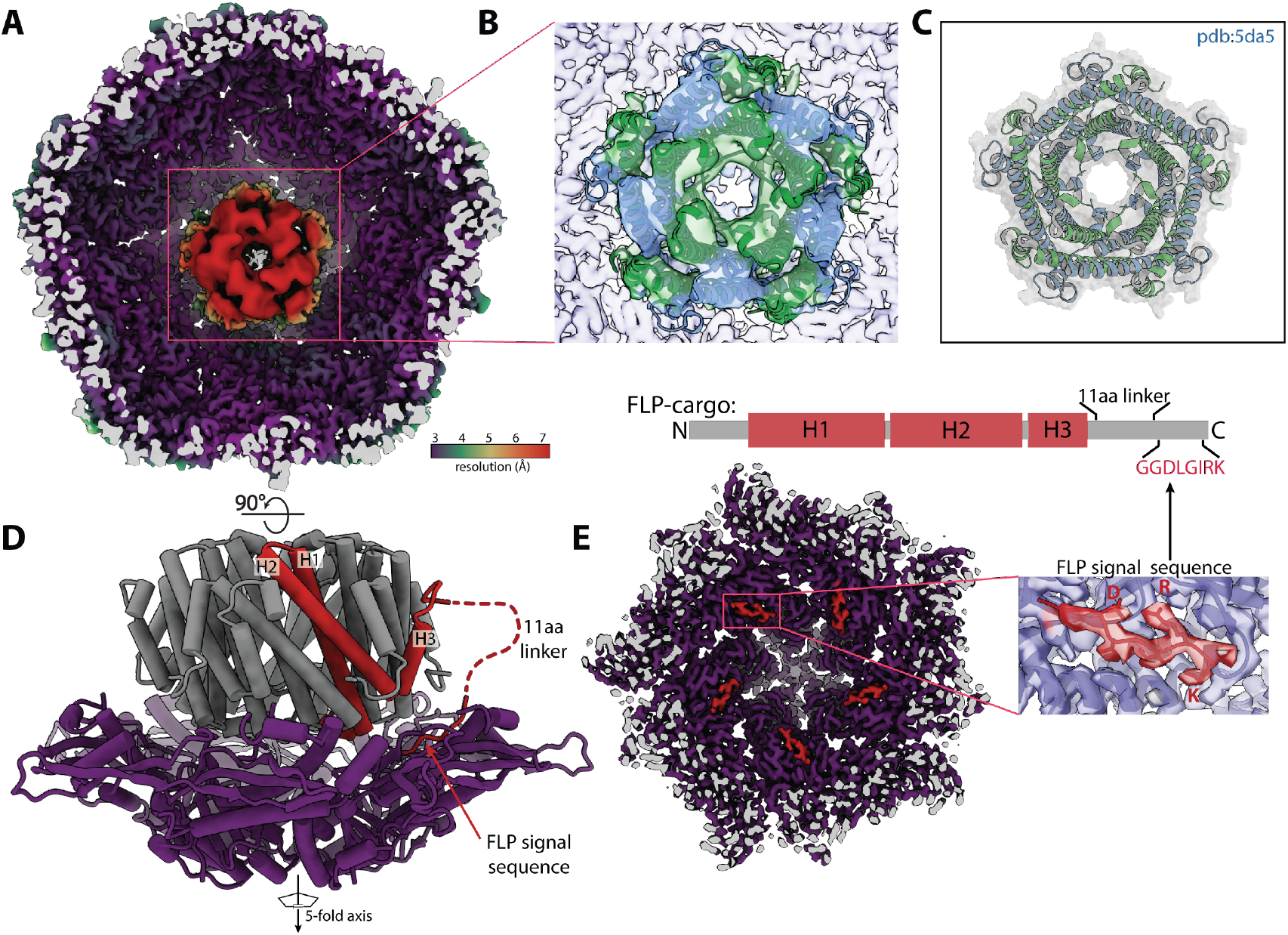
The 5-fold FLP cargo sits on the 5-fold axis of encapsulin. **(A)** Cryo-EM map resulting from symmetry expansion and focused classification, colored by local resolution, with the shell at ∼3.5wÅ resolution and the FLP cargo at ∼7Å. **(B)** Unambiguous docking of a known FLP decamer structure (PDB:5da5), with the crystal structure shown by itself in **(C). (D)** is a 90° rotation and the atomic model for the map shown in **(A)**. This structure shows how the FLP sits at the 5-fold axis as well as the flexible linker that connects the body of the FLP protein to the C-terminal 8 residue signal sequence that connects the FLP to the encapsulin shell **(E)**, confirming what had been identified in previous studies (*4*).

Upon resolving the FLP density, it is clear that the FLP cargo symmetry does indeed match the symmetry axis of the shell. The interactions between the cargo and shell are somewhat flexible, as permitted by the flexible attachment of the known 8 amino acid signal sequence and the main body of the FLP (Figure 4 D,E). The FLP, which is a decamer of heterodimers with 5-fold symmetry, is roughly aligned over the 5-fold axis of the icosahedron. Although this has been intuitively assumed in the previous literature, our structural analysis offers proof of this symmetry matching by providing a structure of both the shell and cargo together. However, due to cargo size and spatial constraints it is clear that not all of the cargo FLPs maintain this symmetry-matched orientation when there are many FLPs inside the encapsulin shell. Steric occlusion appears to distribute the FLP cargos in a more space-filling orientation, while the strongest density persists at one of the pentameric axes of the shell. Based on our finding, we speculate that the 3-fold symmetry of the dye-decolorizing peroxidase (DyP) cargo mentioned briefly in the introduction and found in related encapsulins should bind the 3-fold axis of the shell. However, a concurrent structural study of the *M. smegmatis* DyP encapsulin appears to have the most defined cargo density positioned proximal to the 5-fold axis of the shell rather than the 3-fold axis (*15*). Without an abundance of structural data on diverse encapsulins, it remains unclear what role symmetry plays in the cargo-shell interaction. Nevertheless, most viruses, encapsulins, and related icosahedral structures often use the 5-fold, 2-fold, or 3-fold axis as a pore to regulate the permeability of the compartment (*6*), so localizing cargo proteins to these axes and pores likely has some biological significance.

### The role of FLP cargo and flavin binding within encapsulins

Encapsulins that package FLP do so in order to regulate iron within the cell. Like ferritin, it is thought that these FLP-encapsulins can oxidize soluble ferrous iron and store it within the encapsulin compartment as a mineralized insoluble ferric oxide core (*4, 9, 10*). The oxidation of iron requires the transfer of an electron from iron to an acceptor. Although this acceptor is often molecular oxygen, a flavin bound to the shell could also serve as an electron acceptor. In order to investigate the role flavin may play in the ferroxidase activity of this encapsulin we compared iron storage capabilities in both the WT FLP-encapsulin, and the W90E mutant lacking the flavin. However, iron mineralization assays under aerobic conditions showed no significant difference between the flavin-bound encapsulin and the W90E mutant without flavin (Figure 5 B). Given that *T. maritima* is an anaerobic fermentative chemoorganotrophic organism, we also performed the iron storage assays under anaerobic conditions. There was no significant difference between the encapsulins with or without the flavin in either aerobic or anaerobic conditions tested (Supplemental Figure 5).

**Figure 5.**
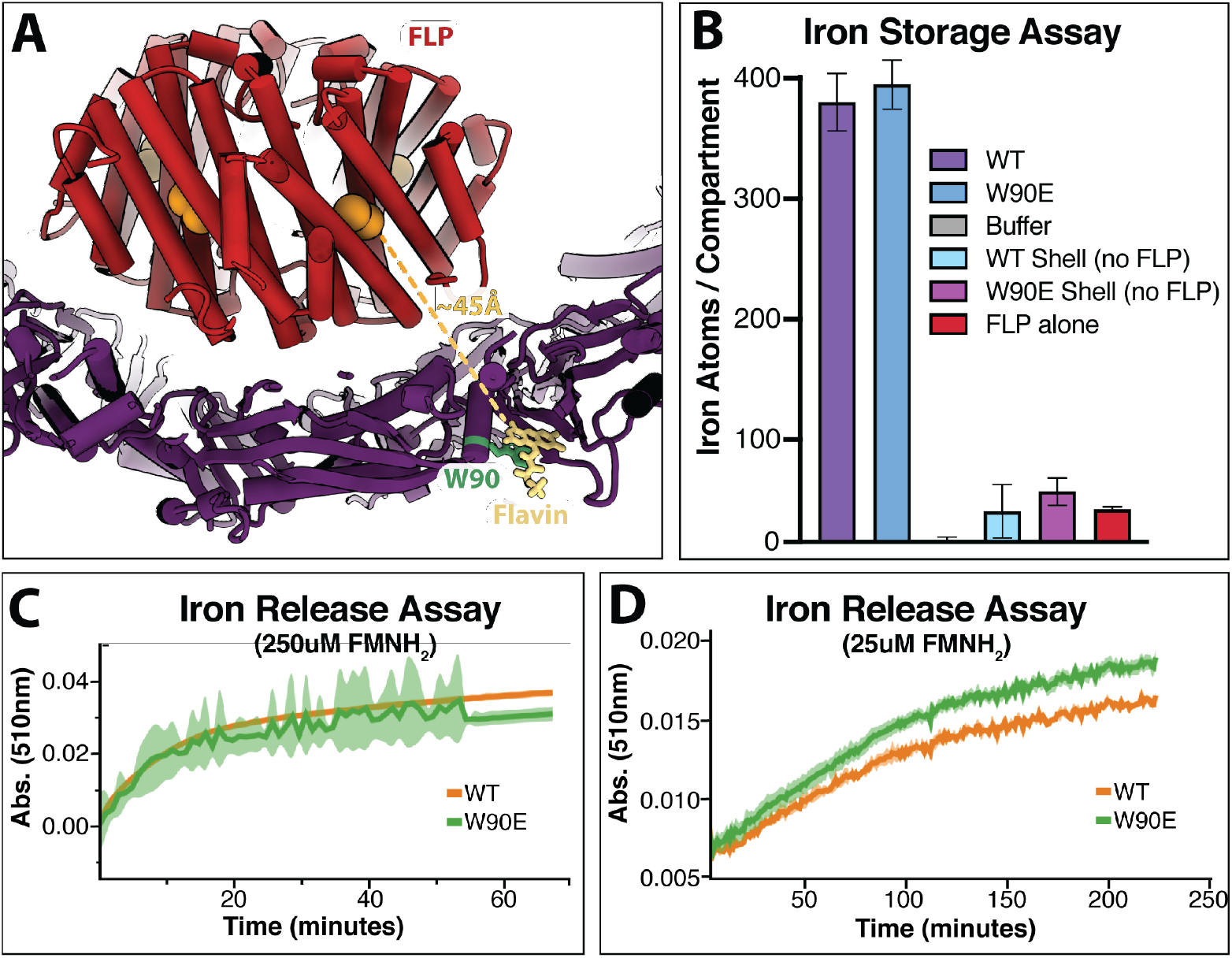
Biochemical data regarding the role of flavin in iron mineralization and iron mobilization. **(A)** Depiction of the distance between the FLP active site and the FMN molecule bound to the shell, with FLP in red, the active site in orange, the shell in purple, the bound FMN in yellow, and the interacting W90 residue in green. **(B)** shows the ferroxidase iron storage activity of various encapsulin constructs. ‘WT’ is the unmodified encapsulin with FLP cargo protein; ‘W90E’ also contains the FLP cargo protein but also represents the W90E mutation to the shell; ‘WT Shell’ does not contain the FLP cargo; ‘W90E Shell’ is also lacking the FLP and contains the W90E mutation; and ‘FLP alone’ represents the FLP cargo protein free in solution that was purified without the encapsulin shell. The opposite activity, the release of iron, is shown in **(C)** at 250uM FMNH2 and **(D)** at 25uM FMNH2. FLP-loaded encapsulin constructs with a ferric oxide core were used for the experiments in **(C)** and **(D)**, with the green curve representing W90E-encapsulin without flavin, and the orange trace is WT FLP-encapsulin. **(C)** and **(D)** lines represent n=3 average, with the envelope showing standard deviation.

Alternatively, the flavin could serve to help mobilize iron if the role of these encapsulins is to release iron when necessary. Under the anaerobic conditions tested, there was also no clear difference between WT-encapsulin and W90E-encapsulin in an iron release assay (Figure 5 C, D). Although these data are negative, the presence of a flavin moiety adjacent to the FLP is suggestive of function and a role may yet be revealed through additional investigation.

A related observation is that the symmetry-expanded structure indicates that the active site of the FLP is ∼45Å away from the shell-bound flavin, on average (Figure 5 A). The symmetry matching and 8 amino acid C-terminal signal sequence that tethers the cargo to the shell likely keeps this distance fairly restrained. However, flexibility may allow for the FLP to occasionally sample distances closer or farther from the shell. Unfortunately, further classification was unsuccessful in resolving classes of cargo in any different orientation. Previous structural data suggests that redox donor and acceptor moieties within a protein are typically within 14Å (*25*). Given our cargo loaded encapsulin structure and this data, it is highly unlikely that this flavin would be the only acceptor involved in iron mineralization. One possibility is that there is a network of aromatics that help shuttle electrons between the flavin and the FLP active site, although the long distances observed in our structure may prohibit this (Supplemental Figure 4). Another possibility is that there is a shuttling molecule between the active site and the shell-bound flavin, as has been shown for other ferritin-like systems (*26*).

Further investigation is necessary to understand the role of this flavin moiety proximal to the FLP cargo for the *T. maritima* encapsulin. One aspect of this system worth consideration is that *T. maritima* is an anaerobic fermentative chemoorganotrophic bacterium. The aerobic conditions used for the iron mineralization assays may allow much higher molecular oxygen as the acceptor molecule than the encapsulin system would otherwise experience naturally. Under cellular anaerobic conditions, the FMN may serve a more pivotal role. In searching for other encapsulins that share the flavin-binding tryptophan residue, almost all of the W90 containing encapsulins came from anaerobic organisms indicating that flavin binding may only be necessary in anaerobic environments. As such, more nuanced experiments are necessary before any claims are made.

Another hypothesis is that there may be an unknown auxiliary protein factor involved in electron transport. For example, the *P. aeruginosa* bacterioferritin systems also contain a protein known as bacterioferritin-associated ferredoxin (*27*). These ferredoxins bind iron-sulfur clusters, are upregulated 200-fold in response to a low iron environment, and are required to liberate any iron from within the bacterioferritin compartments (*27*). A similar undiscovered factor may also play a role within the *T. maritima* FLP-containing encapsulins, which will require further investigation.

## Materials and Methods

### Construction of the T. maritima shell and cargo plasmid

The construct for the *T. maritima* encapsulin shell and FLP-cargo genes was cloned as described previously(*12*) with a few exceptions. The encapsulin shell gene was ordered and codon optimized for *E. coli* from Genewiz. The FLP-cargo gene was ordered from IDT as a gBlock that was codon optimized for *E. coli* expression and included a His_6_-tag for purification. First, the shell gene was cloned into a pET14 vector followed by the FLP-cargo gene upstream with a higher affinity ribosome binding site (Figure 1 A). The rationale for the higher RBS was to maximize the amount of cargo that was expressed and subsequently encapsulated within the shell. For the introduction of the W90E mutation within the shell, golden-gate cloning was performed to introduce the site-specific mutation. All constructs were transformed into DH5α cells for plasmid amplification and sequence verification.

### Protein purification

The purification for all constructs except FLP was adapted from a previous study (*12*). For expression, the pET14 vector containing the encapsulin shell and/or FLP-cargo gene were transformed into BL21(DE3) cells and grown in Terrific Broth media at 37°C. Once the cells grew to and OD_600nm_=0.3, the cells were cooled to 18°C and induced with 500µM IPTG and grown for ∼16hrs. Cells were harvested by centrifugation at 4,000rpm at 4°C for 20min and frozen at −20°C until purified.

The cell pellet was thawed and resuspended in buffer A (50 mM Tris-HCl, pH 7.4, 150 mM NaCl, and 5 mM 2-mercaptoethanol). Resuspended cells were then incubated at room temperature (RT) for 15min in the presence of 100 μg RNase A, 10U DNase, 1mM MgCl_2_, and 2mg lysozyme. Cells were lysed by sonication and the lysate was clarified by centrifugation in a JA-20 rotor at 22,500g for 20min. Following clarification, the supernatant was collected and placed in an 80° C incubator for 3hrs. The supernatant was centrifuged in a JA-20 rotor at 25,000g for 25min to pellet aggregated protein. Following the spin, the supernatant was collected and ammonium sulfate was added to a final concentration of 35%. After a 30min RT incubation, precipitated protein was removed by centrifugation in a JS-5.3 rotor at 6800 g for 20min. The supernatant was collected and ammonium sulfate was again added to reach a final concentration of 75%. The supernatant was incubated for 30min at RT and precipitated proteins were pelleted by centrifugation in a JS-5.3 rotor at 6800 g for 25 minutes. The supernatant was discarded and the pellet was resuspended with buffer A. The resuspended protein was dialyzed into buffer B (20 mM Tris-HCl, pH 7.4, 150 mM NaCl and 5 mM 2-mercaptoethanol) to remove any remaining ammonium sulfate. Dialyzed protein was centrifuged in a type 50.2 Ti rotor at 125,000g for 3hrs to pellet the nanocompartments. The pellet was resuspended in buffer A and loaded onto a Superose 6 10/300 GL column equilibrated with buffer C (50 mM Tris-HCl, pH 7.4 and 150 mM NaCl). Unless otherwise indicated, all Superose 6 columns were run at 0.5 mL/min. Fractions containing assembled encapsulins were collected and concentrated with a Vivaspin 6 (100 kDa MWCO) concentrator. Purified protein was supplemented with glycerol (10% final concentration), flash frozen in liquid N_2_, and stored at −80° C until needed.

FLP was purified by nickel immobilized metal ion affinity chromatography (IMAC). Cell pellets were resuspended in 50 mM Tris-HCl, pH 7.4, 150 mM NaCl, and 10 mM imidazole, and incubated at RT for 15min in the presence of 100 μg RNase A, 10U DNase, 1mM MgCl2, and 2mg lysozyme. Cells were lysed by sonication and the lysate was clarified by centrifugation in a JA-20 rotor at 40,000g for 25min. Cleared lysate was incubated with Ni-nitrilotriacetic acid (NTA) resin from Qiagen for 1hr at RT. The resin was washed with 20 resin volumes of 50 mM Tris-HCl, pH 7.4, 500 mM NaCl, and 10 mM imidazole followed by 20 resin volumes of 50 mM Tris-HCl, pH 7.4, 150 mM NaCl, and 30 mM imidazole. Bound FLP was eluted with 3 resin volumes of 50 mM Tris-HCl, pH 7.4, 150 mM NaCl, and 350 mM imidazole. The Ni-NTA elute was dialyzed into buffer B, concentrated with a Vivaspin 6 (10 kDa MWCO), and loaded onto a Superose 6 column equilibrated with buffer C. Fractions containing FLP were collected and concentrated with a Vivaspin 6 (10 kDa MWCO) concentrator. Purified protein was supplemented with glycerol (10% final concentration), flash frozen in LN2, and stored at −80° C until needed.

### Absorbance and Fluorescence Spectroscopy Measurements

All absorption spectra were acquired in a Cary 50 Bio UV-Visible spectrophotometer. Each purified protein or metabolite was diluted to 20 μM in a buffer containing 150mM NaCl and 50mM Tris-HCl pH 7.4. Samples were placed in a Fisherbrand quartz cuvette and the absorbance was collected from 600nm to 200nm on the slowest scan speed. Fluorescence emission spectra were collected on a Tecan Infinite M1000 PRO spectrophotometer. For native fluorescence, each protein was diluted to 20 μM and each metabolite was diluted to 2 μM in a buffer of 150mM NaCl and 50mM Tris-HCl pH 7.4. Samples were excited at 450 nm with a 5 nm bandwidth. Emission data was collected from 500 nm to 600 nm in 1 nm steps with a 5 nm bandwidth. All samples were measured in triplicate and plotted with Datagraph as the mean value ± standard deviation. For denatured fluorescence, each protein and metabolite were diluted as above (20μM and 2μM, respectively) plus 7M guanidine hydrochloride (GuHCl). Measurements, parameters, and plotting were identical for the both the denatured and native fluorescence samples.

### Mass Spectroscopy

Purified encapsulins were analyzed by LC-MS. The sample was buffer exchanged into water and then injected onto a Waters XBridge C18 column with an Agilent Infinity 1260 HPLC. The sample was eluted with a gradient of water to acetonitrile both supplemented with 0.1% formic acid. Mass analysis was performed with an Agilent 6500 Q-TOF operated in positive ion mode. The organic solvent denatures the encapsulin particles on the column thus releases the bound flavin species.

### Cryo-EM sample preparation and data collection

Samples were prepared on Protochips CFlat 1.2/1.3-T grids. Grids were glow discharged on a Tergeo Plasma cleaner prior to use. A frozen aliquot of the *T. maritima* encapsulin was diluted to 2mg/mL in the same buffer as used for size exclusion (20mM NaPO_4_ pH 7.5, 50mM NaCl) in order to remove the 10% glycerol cryoprotectant used during freezing the aliquots. The 2mg/mL sample was supplemented with 0.05% NP-40 to aid in ice uniformity throughout the cryo-EM grid. 4uLs of sample were applied to the grid, and immediately plunge-frozen in liquid ethane using a Vitrobot Mark IV (blot force 10, 2 sec blot, 100% humidity 4°C). The grid was then loaded into a Gatan 626 side entry holder and inserted into a low-base Titan microscope operating at 300 keV. 1,265 micrographs were collected using a Gatan K2 in counting mode at a pixel size of 0.82Å and a total electron dose of 60e^-^/Å^2^.

### Data processing and atomic model refinement

Data was processed within the RELION/3.1 pipeline (*28*). Individual frames were aligned with MotionCorr2 (*29*) and defocus estimation was performed using CTFFIND4 (*30*). Roughly 1,000 particles were manually picked and 2D classification was performed to generate templates for auto-picking within RELION. Initially 86,124 particles were extracted. Two rounds of 2D classification were performed, followed by a round of 3D classification resulting in 38,952 particles in the final reconstruction, which was performed with I1 symmetry. The reconstruction was subjected to multiple iterations of CTF refinement and Bayesian polishing within RELION (*31*), ultimately yielding a high-resolution structure for the shell (*31*). The previously determined shell structure (pdb:3DKT) was used as an initial model, and phenix.real_space_refine (*32*) was used to refine the coordinates of 1 subunit surrounded by its nearest neighbors and with FMN bound. The single ASU was used for the model for the FSC_model-vs-map_, and Chimera was used to create the icosahedral biological matrix. Unfortunately, the symmetrization averaged out the sub-stoichiometric cargo density and therefore did not allow for visualization of the FLP. A C1 reconstruction revealed modest density for the FLP (Supplemental Figure 3). Signal subtraction and re-refinement improved the FLP cargo density only slightly. Therefore, symmetry expansion and focused classification were performed. The process yielded a much-improved density for the FLP cargo protein where we were able to unambiguously rigid-body dock the FLP crystal structure (pdb:5DA5) (*10*).

### Ferroxidase (Iron Storage) Assays

Purified protein was diluted in a buffer of 50mM Acetate, pH 6.0 and 30 mM NaCl. FLP loaded WT encapsulin and FLP loaded W90E encapsulin were diluted to 20μM of FLP. Ammonium ferrous sulfate (2.5mM) was prepared anaerobically in 0.1% HCl. The ferrous iron was diluted ten-fold into the protein, mixed, and then immediately loaded into a Cary 50 Bio UV-Visible spectrophotometer. Absorbance at 310nm was collected every second for 30 minutes. To normalize between all the samples, the initial 310nm absorbance value was set as the 0-absorbance value. Data was plotted using Datagraph.

### Iron Release Assays

Reduced FMN, FMNH_2_, was generated by mixing FMN and NADH anaerobically and incubating overnight. Purified protein was diluted in a buffer of 150mM NaCl and 50mM Tris-HCl pH 7.4 and made anaerobic by purging with a H_2_:N_2_ gas mixture. Anaerobic 1,10 phenanthroline (750μM final concentration) and FMNH_2_ (25μM or 250μM) were added to the sample. Iron release was monitored by the gain in absorbance at 510nm in a Tecan M200 plate reader.

## Acknowledgements

We would like to thank the Coates lab for help with the anaerobic iron release assays. We would also like to thank Patricia Grob for EM support and Abhiram Chintangal for computer support. Thanks to B.G. for intellectual insights. B.L. was supported by NSF-GRFP (1106400). E.N. is a Howard Hughes Medical Institute investigator and D.F.S. was supported by a grant from the U.S. Department of Energy (DE-SC00016240).

## Author Contributions

BL, CCA, and DFS designed research; BL, CCA, RJN, and LMO performed research; BL, CCA, RJN, LMO, EN, and DFS analyzed data, BL, CCA, and DFS wrote the paper with assistance from RJN, LMO, and EN.

## Supplemental Information

**Supplemental Figure 1.**
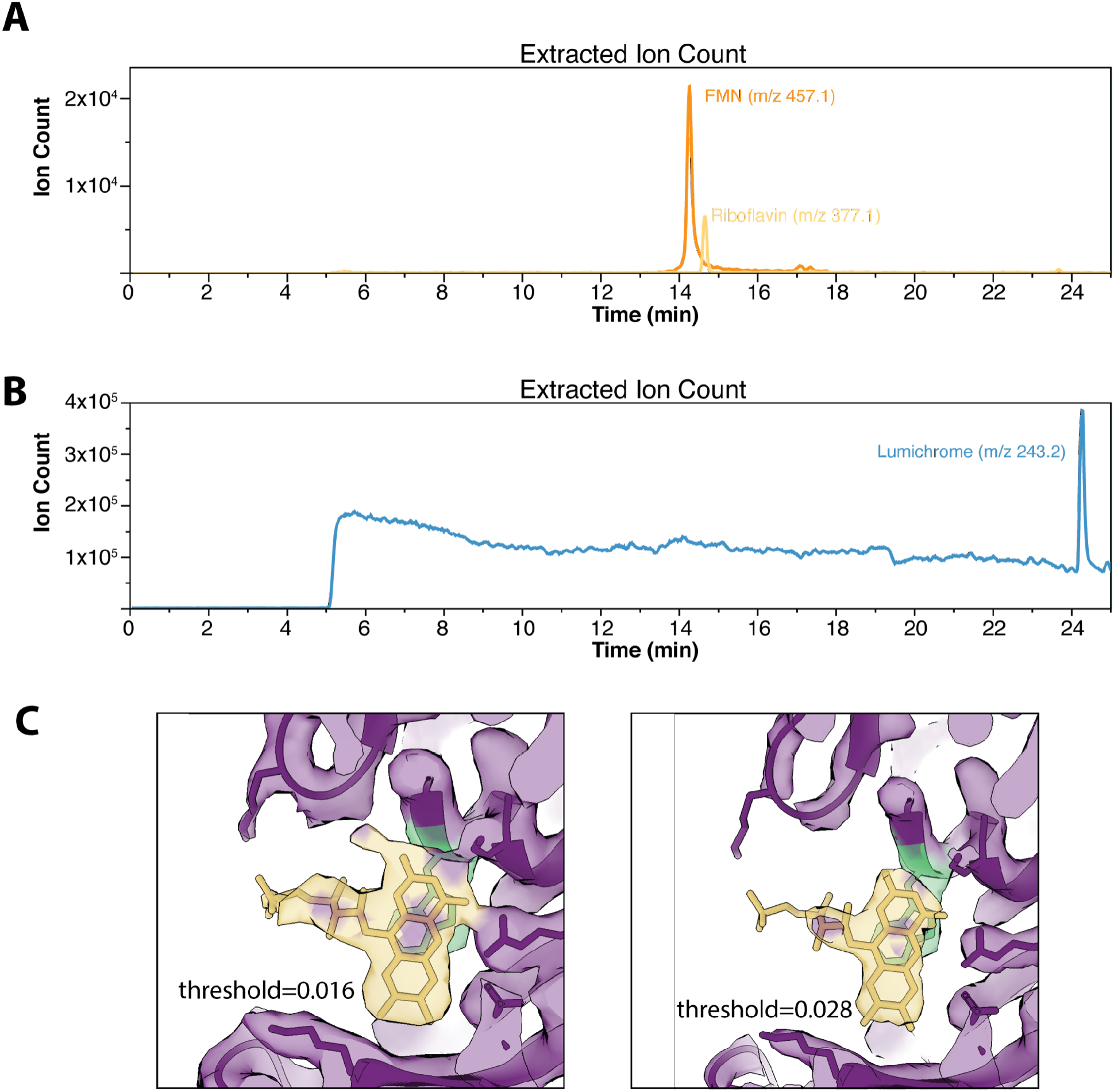
Mass spectrometry confirms flavin presence in the *T. maritima* encapsulin. **(A)** and **(B)** Extracted ion counts show that FMN (orange) riboflavin (yellow), and lumichrome (blue) are all present within a pure *T. maritima* encapsulin sample. **(C)** CryoEM density of the flavin at two different thresholds showing density for the FMN and riboflavin R-group that is absent in the lumichrome degradation product.

**Supplemental Figure 2.**
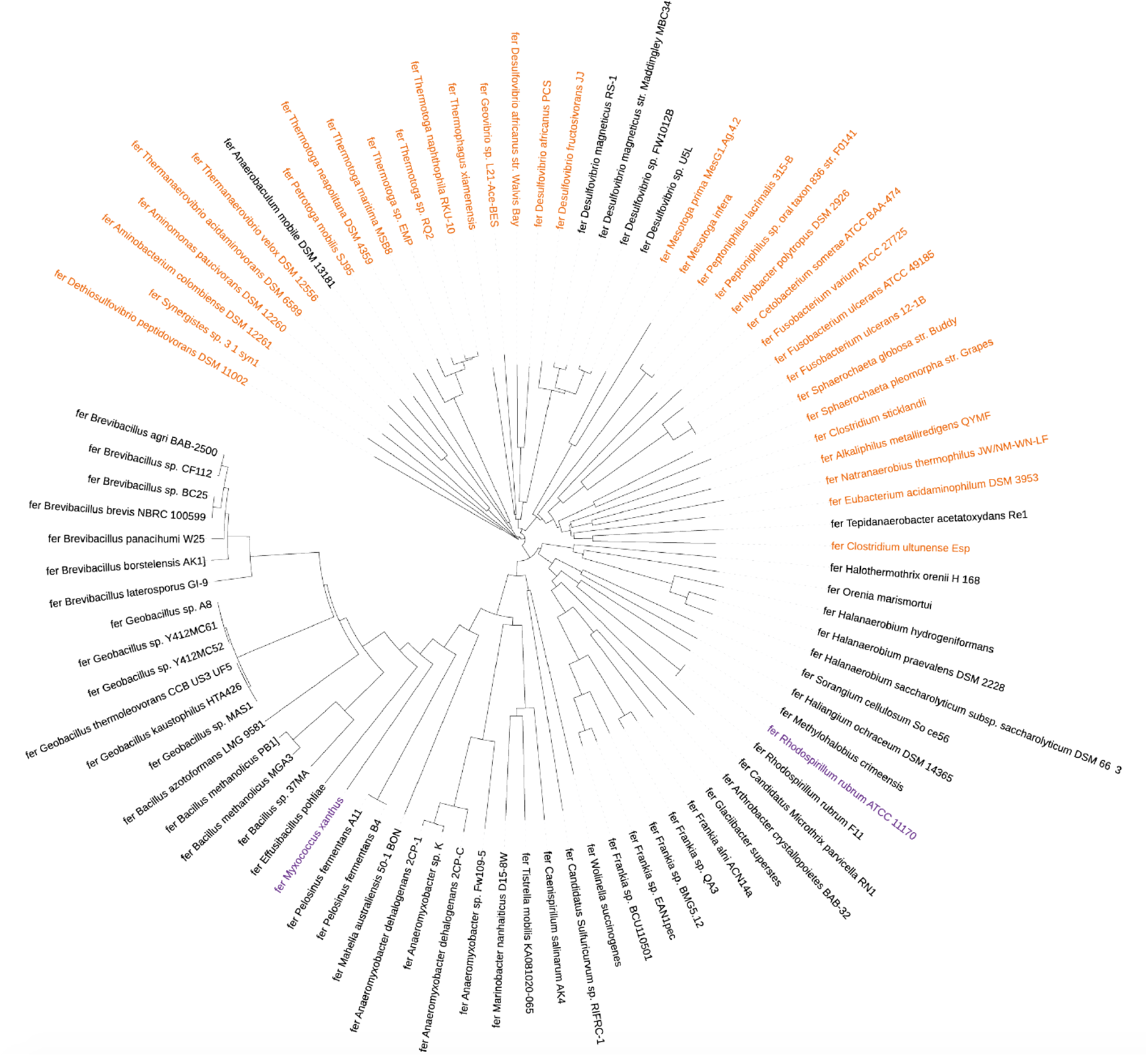
Phylogenetic tree of encapsulins with FLP-cargo. Encapsulins with a tryptophan at position 90 are colored in orange. Previously characterized encapsulins are colored in purple.

**Supplemental Figure 3.**
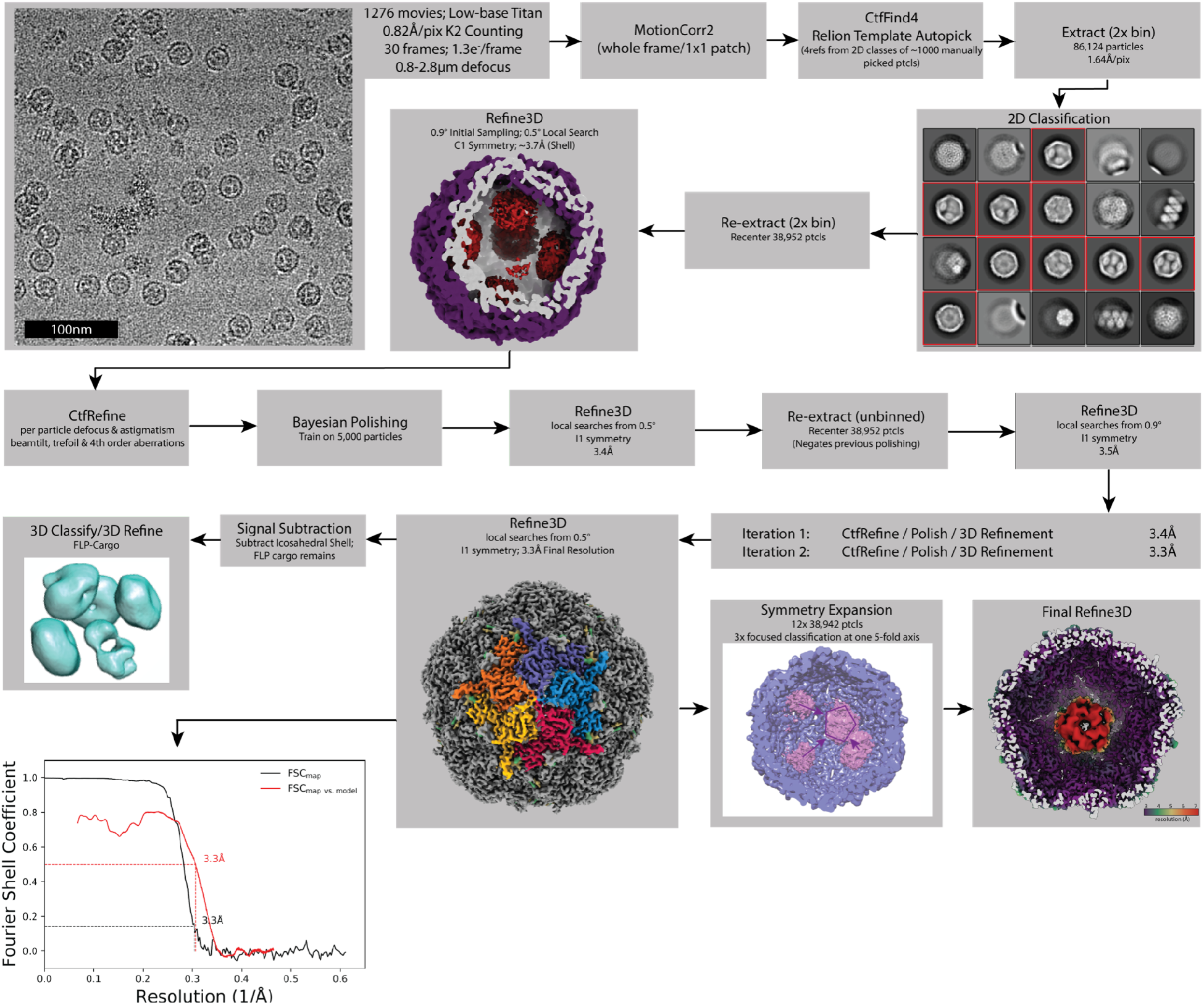
CryoEM Processing Pipeline. Workflow used for obtaining a high-resolution encapsulin shell, as well as the symmetry expansion technique used for defining one of the possible FLP-cargo orientations relative to the shell.

**Supplemental Figure 4.**
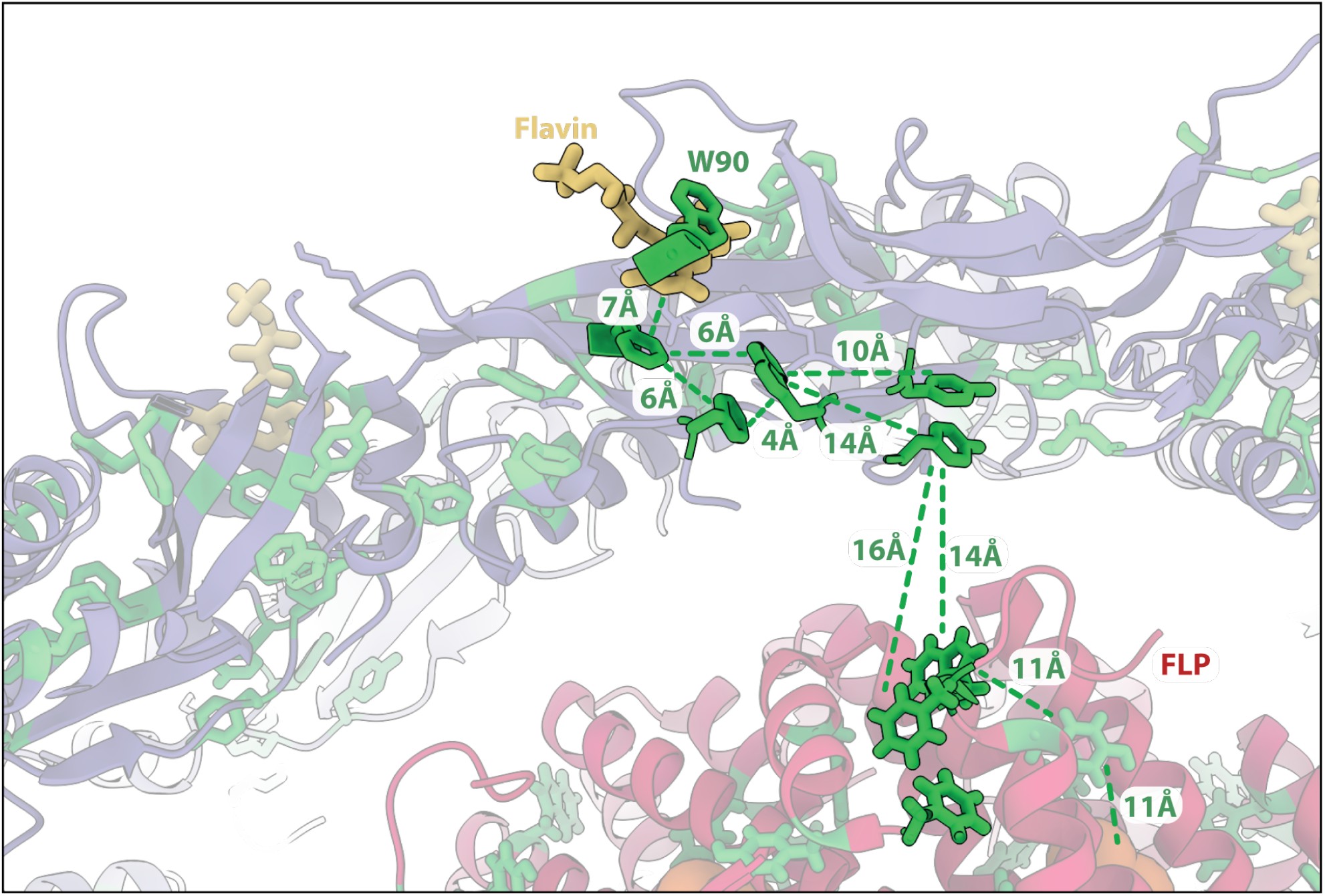
Aromatics positioned between the flavin molecule and the FLP active site. Distances (in angstroms) between aromatic side chains that could serve as an electron conduit between the flavin (yellow) on the shell periphery and the active site of the encapsulated FLP cargo (orange)

**Supplemental Figure 5.**
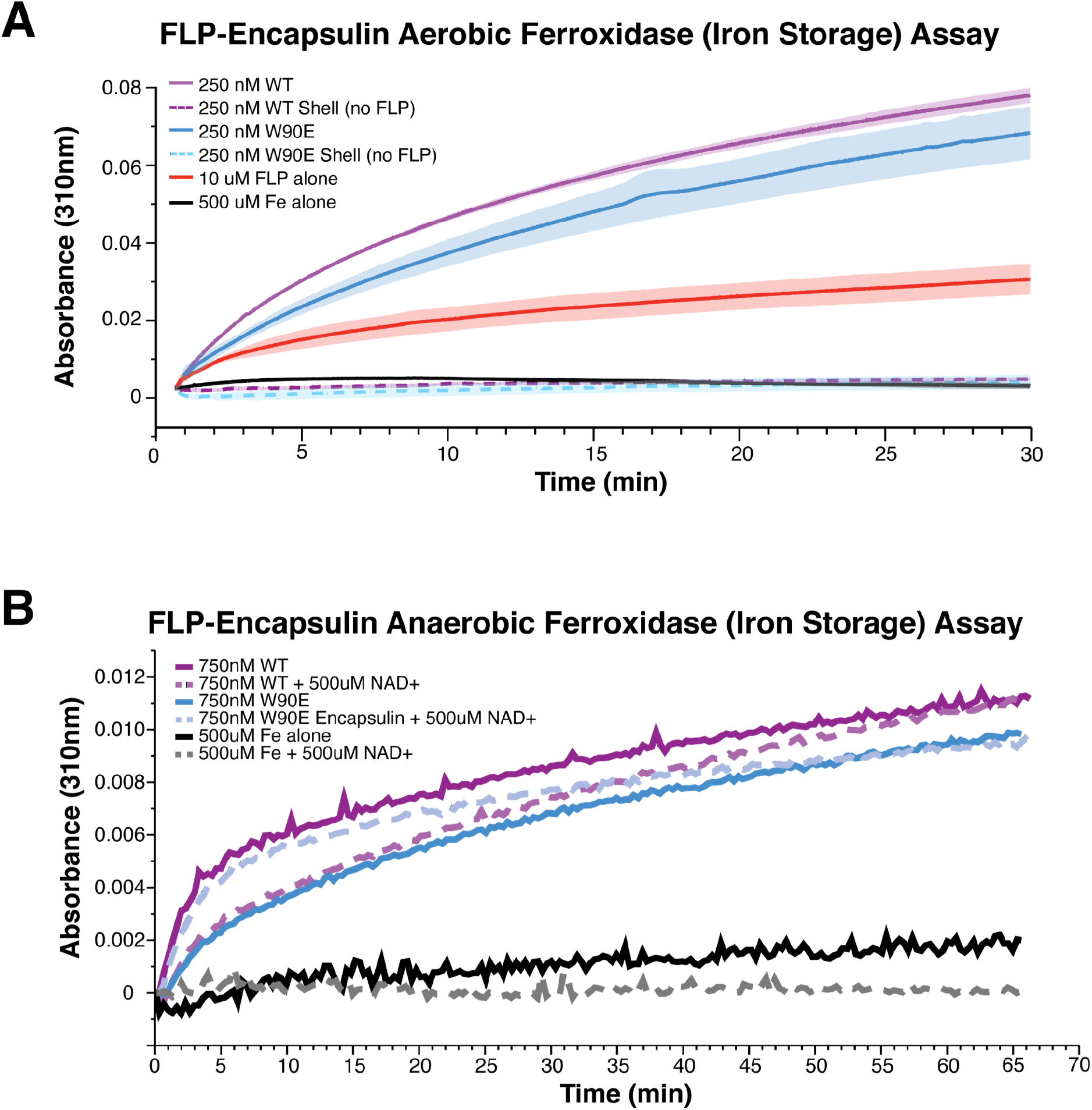
The ferroxidase activity of different encapsulin constructs. All assays were performed in the presence of 500uM Fe. **(A)** Shows the ability of different encapsulins to store iron under aerobic conditions. Traces are n=3 average with the envelope representing STD. **(B)** Ferroxidase activity under anaerobic conditions. Traces represent an average of n=3. Both WT encapsulin and the W90E mutant retain the ability to store iron regardless of aerobic/anaerobic, with no significant statistical difference between the two.

**Supplemental Table 1:**
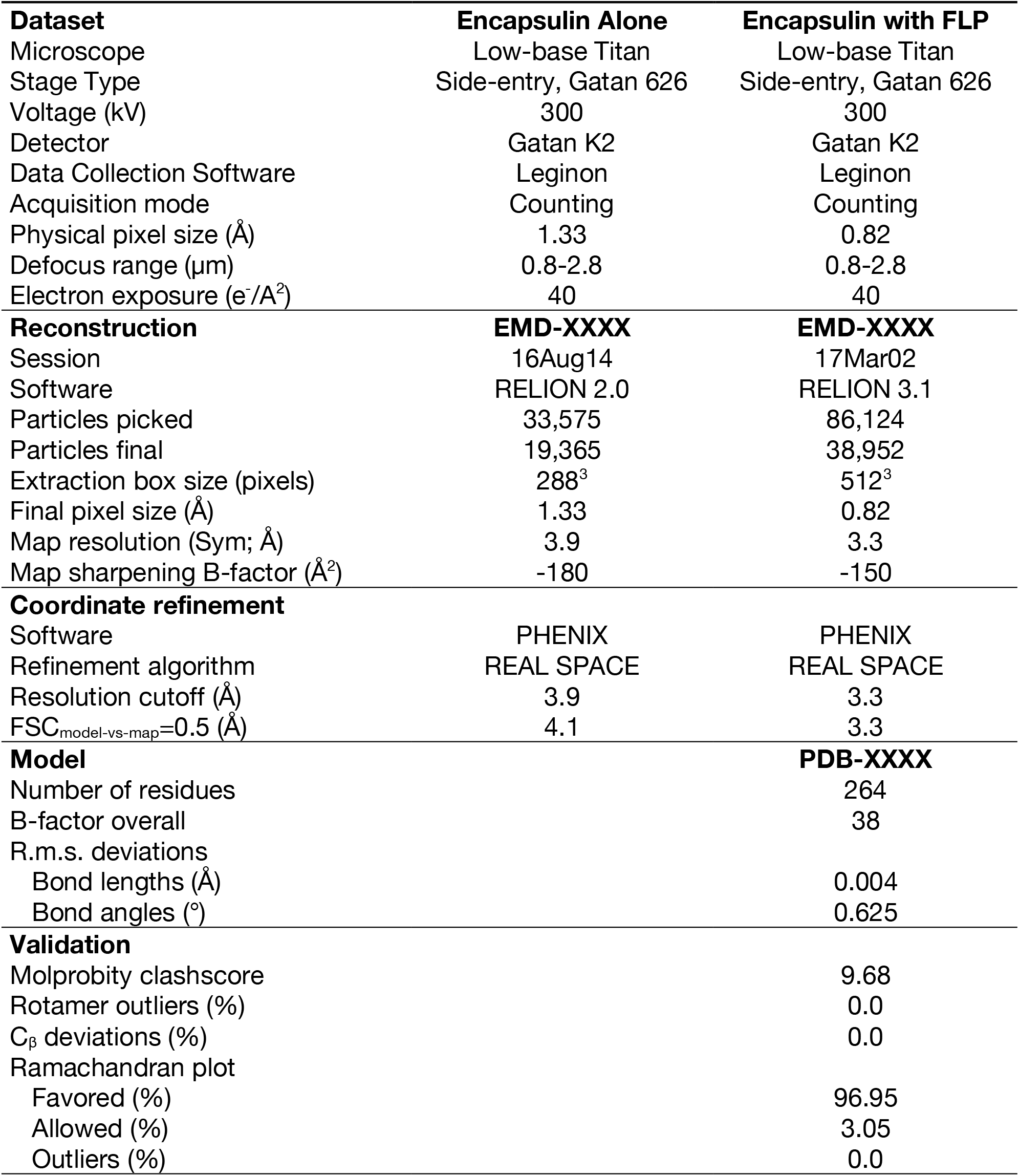
Data collection, 3D reconstruction, and refinement statistics.

